# Generation, characterization and exploiting caprine herpesvirus 1 secreted glycoprotein D

**DOI:** 10.1101/2025.06.30.662481

**Authors:** Sergio Minesso, Amienwanlen Eugene Odigie, Valentina Franceschi, Simone Taddei, Vittorio Madia, Maria Tempesta, Gaetano Donofrio

**Affiliations:** Department of Veterinary Science, University of Parma, 43126, Parma, Italy; Department of Veterinary Medicine, University of Bari, 70010 Valenzano, Italy

## Abstract

Caprine herpesvirus 1 (CpHV-1), a member of the *Herpesvirales* order, *Herpesviridae* family, *Alphaherpesvirinae* subfamily, and *Simplexvirus* genus, is classically associated to two distinct clinical syndromes. In kids, CpHV-1 induces severe systemic disease with high morbidity and mortality, in adult goats, the infection leads to genital lesions such as vulvovaginitis or balanoposthitis, with abortions occurring mainly in the second half of gestation. CpHV-1 shares several biological characteristics with human herpesvirus 2 (HSV-2), including molecular features, tropism for vaginal epithelium, genital lesion nature and latency in the sacral ganglia. These features make CpHV-1-infected goats a reliable animal model for studying human herpesvirus-induced genital disease, employable for pathogenic research, as well as the development of new vaccines and antiviral agents. Recent full sequencing of CpHV-1 genome has identified at least ten genes encoding glycoproteins. Among these, glycoprotein D (gD) has been characterized but not yet exploited for immunogenic or diagnostic purposes. In this study, the structural features of CpHV-1 gD were predicted using in silico analysis. A truncated version of gD lacking the transmembrane domain (Sec-gD) was subsequently generated and expressed in mammalian cells, enabling its secretion into the culture medium. Despite the structural modifications, Sec-gD retained a conserved glycosylation pattern, as confirmed by PNGase F treatment. Furthermore, the antigenic properties of Sec-gD were preserved, as demonstrated by reverse serum neutralization assays. Notably, the culture supernatant containing Sec-gD was directly usable in diagnostic enzyme-linked immunosorbent assays, supporting its potential as a valuable tool for both diagnostic and immunization strategies.

**Importance:** CpHV-1-infected goats represent a large animal model for studying human herpesvirus-induced genital disease, and could be utilized for pathogenic research, as well as for the development of new vaccines and antiviral agents. CpHV-1 gD can be efficiently produced and rescued from the supernatant of transfected mammalian cells, retaining its immunogenic properties and could be employed for immunogenic and diagnostic purposes.

## Introduction

Caprine herpesvirus 1 (CpHV-1) is a member of the Herpesvirales order, within the Herpesviridae family, Alphaherpesvirinae subfamily, and Simplexvirus genus (Davison, 2010). This virus is associated with two distinct clinical syndromes in goats: a fatal systemic illness in kids (Van der Lugt and Randles, 1993), and a genital disease in adults, which can manifest as balanoposthitis (Tarigan et al., 1987), vulvovaginitis (Tempesta et al., 1999a), and abortion (Keuser et al., 2002). Although only one complete genome sequence of CpHV-1 has been determined so far (https://www.ncbi.nlm.nih.gov/nuccore/NC_076509; Accession number: NC_076509), restriction site maps have been developed using double digestion and cross-hybridization of individual restriction fragments. The molecular weight of CpHV-1 DNA, estimated by summing the weights of fragments generated by various endonucleases, is approximately 137 kbp (Engels et al., 1987; Hao et al., 2020). From a pathogenic perspective, CpHV-1 infection typically begins at the respiratory or genital mucosa. The virus then disseminates systemically via a mononuclear cell-associated viremia, potentially leading to abortion in pregnant animals. CpHV-1 is shed through ocular, nasal, and genital secretions, with the genital tract considered the primary site for viral entry and persistence within herds (Tempesta et al., 1999a). In kids, CpHV-1 causes a severe systemic disease marked by high morbidity and mortality, with ulcerative and necrotic lesions throughout the gastrointestinal tract. In adult goats, the infection results in genital lesions such as vulvovaginitis or balanoposthitis. Abortions typically occur in the second half of gestation and can be experimentally induced through intranasal or intravenous inoculation of pregnant goats (Uzal et al., 2004). Following intravaginal infection, the virus establishes latency in the sacral ganglia. Reactivation may occur under physiological stress, particularly during the breeding season, and may be influenced by hormonal changes during estrus. However, experimental reactivation is challenging and generally requires high doses of dexamethasone (Tempesta et al., 1999b). Notably, CpHV-1 shares several biological characteristics with human herpesvirus 2 (HSV-2) and bovine herpesvirus 1 (BoHV-1), including molecular features, tropism for vaginal epithelium, the nature of genital lesions, and latency in the sacral ganglia (Camero et al., 2010). Caprine herpesvirus 1 (CpHV-1) genomes have been recently full sequenced (https://www.ncbi.nlm.nih.gov/nuccore/NC_076509) (Accession number: NC_076509), exhibit the characteristic structure of a class D herpesvirus genome. This genome class is composed of a linear double-stranded DNA molecule, organized into a unique long (UL) segment and a unique short (US) segment. The US segment is flanked by internal and terminal inverted repeat sequences (IR and TR, respectively). The complete genome sequence of CpHV-1 has revealed the presence of at least ten genes encoding glycoproteins. These glycoproteins have been identified and named based on their homology with those of herpes simplex virus type 1 (HSV-1) (Schwyzer and Ackermann, 1996). Among these glycoproteins, glycoprotein D has been characterized (Keuser et al., 2006) however, not exploited as an immunogen or in terms of diagnostics. In this work, we successfully expressed and characterized gD in a secreted form (sec-gD) in the medium of a mammalian cell and the medium containing sec-gD could be directly employed for the development of a diagnostic enzyme-linked immunosorbent assay or immunization purposes.

## Materials and Methods

### *In sicilo* analysis of CpHV-1 glycoprotein D

The amino acid sequence of CpHV-1 glycoprotein D (gD) was retrieved from NCBI (https://www.ncbi.nlm.nih.gov/protein/; accession no. AAZ66865.1) in FASTA format and used as input for the downstream analyses. PSIPRED 4.0 (McGuffin et al., 2000) and GOR IV web server (Garnier et al., 1996) were employed for secondary structure prediction. Prediction of transmembrane helices and signal peptide was performed with Phobius web server (Kall et al., 2007); visualization of protein topology was carried out with Protter v. 1.0 (Omasits et al., 2014). Prediction of glycosylation sites was performed with NetGlyc 1.0. (Gupta and Brunak, 2002). Template-based 3D modelling of CpHV-1 gD was performed using the SWISS-MODEL web server (Waterhouse et al., 2018). A total of 172 templates matched the CpHV-1 gD sequence. Due to the unavailability of CpHV-1 gD templates, the prediction was based on crystal structures of the glycoprotein D from related herpesviruses **(Supplementary File 3)**. From these, the 10 most suitable templates were selected for further modelling. The resulting best-fitting model was selected based on sequence coverage, identity, Global Model Quality Estimate (GMQE) and the QMEANDisCo global value. GalaxyRefine (Ko et al., 2012) was then employed to enhance the prediction accuracy. Both crude and refined models were evaluated with the SWISS-Model structure assessment tool.

### Cells

HEK 293 T (human embryo kidney cells; ATCC: CRL-11268), MDBK (Mardin Darby Bovine Kidney; ATCC: CRL-6071) and BEK (bovine embryo kidney; Istituto Zooprofilattico Sperimentale, Brescia, Italy; BS CL-94) were cultured in complete Eagle’s minimal essential medium (cEMEM). cMEM was supplemented with 2 mM of L-glutamine, 1 mM of sodium pyruvate, 100 IU/mL of penicillin, 100 μg/mL of streptomycin, 0.25 μg/mL of amphotericin B) and 10% FBS. Cells were cultured in a humidified incubator at 37 °C/5% CO_2_. All the supplements for the culture medium were purchased from (Gibco, Segrate (MI), Italy).

### Construct generation

CpHV-1 gD secreted fragment (Sec-gD), lacking the trans-membrane domain, was obtained by PCR amplification from pINT2-CMV-gDcpgD_106_ (Donofrio et al., 2013) with NheI-Cap-gD sense (5′-CCCGCTAGCATGTGGGCCCTCGTGCTCGCAGCGCTAAGC-3′) and BamHI-HA-Cap-gD antisense (5’-GGGGGATCCTTAGGCGTAATCGGGCACGTCGTAGGGGTACGGCGCGGCGGGCGGGAGGGTAGGC-3′) primers. This pair of primers introduced a NheI restriction site at the 5’ and an HA tag of the ORF, followed by a BamHI site at the 3’terminal. The PCR amplification reaction was implemented in a final volume of 50 μl, containing 20 mM Tris–hydrochloride pH 8.8, 2 mM MgSO4, 10 mM KCl, 10 mM (NH4)2SO4, 0.1 mg/ml BSA, 0.1% (v/v) Triton X-100, 5% dimethyl sulfoxide (DMSO), 0.2 mM deoxynucleotide triphosphate, and 0.25 μM of each primer. 1U of Pfu recombinant DNA polymerase (Thermo Fisher Scientific), was used to amplify 100 ng of template DNA over 35 repeated cycles, including 1 min of denaturation at 94°C, 1 min of annealing at 60°C, and 1 min of elongation at 72°C. The resulting Sec-gD-HA amplicon was restriction digested with NheI/BamHI and subcloned in pEGFP-C1 (Clontech), previously digested with the same enzymes, to generate the pCMV-Sec-gD construct.

### Transient transfection

HEK 293 T cells were plated into 175cm^2^ flasks (5 × 10^6^ cells/flask) and incubated at 37°C with 5% CO^2^.At sub-confluent density, the culture medium was removed, and the cells were transfected with pCMV-Sec-gD or pEGFP-C1 (as a mock control; Clontech) using Polyethylenimine (PEI) transfection reagent (Polysciences, Inc.). Briefly, DNA was mixed with PEI in a ratio 1:2.5 DNA:PEI in 3.500 ml of serum-free Dulbecco’s modified essential medium (DMEM) with high glucose (Euroclone) and incubated 15 min at room temperature. Next, 4x volumes of serum-free medium were added, and the transfection solution was transferred onto the cells monolayer and left for 6 hs at 37°C with 5% CO2, in a humidified incubator. The transfection mixture was then replaced with 21 ml of DMEM/F12 (1:1) and incubated for 48 hs. The cell supernatants, containing Sec-gD protein, were then harvested, clarified at 2500rpm at 4°C and stored at -80°C.

### Immunoblotting

Different amounts of Sec-gD protein supernatants samples were electrophoresed on 10% SDS-PAGE after total protein quantification with a BCA Protein Assay kit (Pierce™, ThermoScientific) and then transferred to PVDF membranes by electroblotting (Millipore, Merck, Rahway, NJ, USA). The membrane was subsequently blocked in 5% skim milk (BD), incubated 1 hour at RT with a primary mouse monoclonal antibody anti-HA tag (G036, Abm Inc., New York, NY, USA), diluted at 1:15,000, and then probed with horseradish peroxidase-labeled anti-mouse immunoglobulin (A9044, Sigma-Aldrich (Merck), Tokyo, Japan), diluted at 1:15,000, and visualized using enhanced chemiluminescence (Clarity Max Western ECL substrate, Bio-Rad, Hercules, CA, USA).

### PNGase F digestion

PNGase F, purchased from NEW ENGLAND BioLabs, was employed following manufacturer’s instructions.. Sec-gD protein-containing supernatants were collected from pCMV-Sec-gD transiently transfected HEK cells after 48 hours of secretion. The samples were then treated with PNGase F, which cleaves between the innermost N-acetylglucosamine (GlcNAc) and asparagine residues from N-linked glycoproteins. PNGase F treated samples were subsequently analyzed by Western immunoblotting as described above.

### Virus Titration

BA-1 strain of CpHV-1, isolated from a latently infected goat, was cultured and titrated in MDBK cell (Tempesta et al., 1999a). The stock viral titre of 10^6.25^ 50% tissue culture infectious doses (TCID_50_)/50 µL was stored at -80^0^C and used for the experiments. Briefly, stock virus was serial 10-fold diluted and inoculated in quadruplicate onto MDBK cells in 96-wells microtiters plates and incubated at 37^0^C in a 5% CO_2_ atmosphere environment. The result was read after 3 days of incubation and viral titres were expressed as logarithmic units calculated by the Reed-Muench method (Tempesta et al., 2007).

### Animals and Experimental Infection

The experimental protocol for goats’ infection was duly authorized (code 48E68, min aut. 869/15.11.2021) and was conducted at the authorized University of Bari experimental animal facility (authorization nr. 06/2023-UT).

Ten five-year-old female goats without neutralizing antibodies to CpHV-1, as demonstrated by seroneutralization assay (SNA), were used in this study. Prior to experimentation, the goats were held under controlled environmental conditions and examined daily for clinical evidence of disease. Nasal and vaginal swabs of each were at the time the goats arrived at the laboratory, and just before inoculation to ensure the absence of ongoing infection.

Goats were intravaginally infected with Ba-1 strain of CpHV-1, each receiving 3.0 mL (2 x 10^6^TCID_50_ per mL) of the virus preparation, and were daily examined for clinical evidence of infection. Observed clinical signs such as hyperemia, oedema, lesions and pain were scored as 0 (absent), 1(mild), 2 (moderate), and 3 (severe). Temperature elevations above normal (38.2 – 38.6^°^C) were graded as 1 (>0.5 – 1 ^°^C); 2 (1.1 – 1.5 ^°^C); and 3 (> 1.5 ^°^C). The score for each animal were monitored and reported daily. Blood samples were taken at day 0-, and 42-days post infection to evaluate antibody response to CpHV-1 by means of SNA. Vaginal swabs were also obtained daily for 11 days post infection to evaluate virus shedding.

### Serum neutralization test

The serum neutralization assay, yielding the highest degree of specificity of all the serological tests, was used to assess seroconversion following procedure described elsewhere (Tempesta et al., 2000). In brief, sera were obtained from goats by venipuncture in EDTA-free vacutainer and heat inactivated at 56 C for 30 minutes. Subsequently, serial two-fold dilutions of each serum from 1:2 up to 1:32 were mixed with 100 TCID50 of BA-1 strain of CpHV-1 in a 96-well microtiter plates. The plates were held for 45 minutes at room temperature, and then, 20,000 MDBK cells in a volume of 0.05 mL of DMEM were added to each well and the plates incubated for 3 to 5 days at 37 C in a 5% CO2 humidified chamber environment. Readings were made when CPE were complete in the virus control cultures and the toter of each serum was expressed as the highest dilution neutralizing the virus in the well.

### Samples collection and ELISA procedure

96 wells microplates (MICROLON HIGH BINDING) were coated overnight at 4°C with 50 ng/well of Sec-gD protein supernatant diluted in 0.1 M carbonate/bicarbonate buffer at pH 9.6. After blocking with 1% bovine serum albumin (BSA; Sigma Aldrich by Merck), goat serum samples at different two-folds dilutions (1/10, 1/20, 1/40, 1/80, 1:160, 1:320, 1:640 and 1/1280) were incubated for 1 hour at room temperature. Serum samples were diluted in DMEM/F12 without serum, collected from HEK293 grown for 48 hours. After three washing steps in Phosphate Buffer Saline (PBS), 50 μL of donkey anti-goat IgG-HRP (SANTA CRUZ BIOTECNOLOGY, Germany) diluted 1:5.000 was added to each well and the plate was incubated as above. Following the final washing step, the reaction was developed with 3,3′,5,5′-tetramethylbenzidine (TMB; Merck), stopped with 0.2 M H_2_SO_4_ and read at 450 nm.

### Serum-neutralization inhibition test

Three heat-inactivated caprine sera previously confirmed to be positive in SNAs against CpHV-1 were selected. Twenty-five microliters of each CpHV-1 neutralizing serum sample were added to the first row of 96-well plates. An equal volume of cMEM (without FBS) was added to each well, and for each serum tested, serial twofold dilutions were performed. Next, 25 μl of medium containing Sec-gD protein were added to each well. Each serum was tested in the presence (+Sec-gD) or in the absence of Sec-gD (−SecgD). Positive and negative virus controls were also included. After 1h of incubation at room temperature, 25 μl of virus suspension containing 100 TCID_50_ (50% tissue cell infectious doses) of CpHV-1 strain Ba.1 (Buonavoglia et al., 1996) were added to each well. After 1 h of incubation at 37°C, 50 µl of a BEK cell suspension (2*10^5^ cells/ml) was added to each well and the plates were incubated for 2 days at 37 °C/5% CO_2_. Expression of viral infectivity and serum neutralizing activity through CPE were detected by microscopy and or by classical crystal violet staining of the cell monolayer. The neutralization antibody titer was expressed as the reciprocal (log 2) of the final dilution of serum that completely inhibited viral infectivity.

### Receiver operating characteristic (ROC) analysis

ROC analysis was carried out using SPSS for Windows (version 29.0.1.0, SPSS Inc., Chicago, USA) and the results were plotted using GraphPad Prism (version 8.0.1, GraphPad Software Inc., Boston, USA).

## Results

### Generation and expression of CpHV-1 gD as a secreted peptide

Based on its amino acid sequence and predictions from Phobius/Protter (http://phobius.sbc.su.se/), which are consistent with findings by Keuser et al. (Keuser et al., 2006), the CpHV-1 gD open reading frame (ORF) is 1224 nucleotides long and encodes a 407-amino acid protein. This protein has a predicted molecular mass of 42.6 kDa and includes a signal peptide (amino acids 1–17), a hydrophobic transmembrane domain (amino acids 360–376) near the C-terminus, and a 31-amino acid cytoplasmic tail **(Fig. 1A** **and Supplementary File 1)**. The full-length gD can be expressed in eukaryotic systems as a membrane-bound protein (Donofrio et al., 2013). Based on this structure, removal of the transmembrane domain was predicted to yield a secreted form of gD **(Fig. 1B** **and Supplementary File 1)**. For achieving this, the extracellular domain of gD was amplified by PCR using primers, with the antisense primer incorporating an HA tag sequence in-frame with the amino terminal rest of protein **(Supplementary File 1)**. The resulting ORF was cloned into an expression vector, generating pCMV-Sec-gD. This construct includes the CMV promoter, the gD ORF lacking the transmembrane domain fused to an HA tag, and the bovine growth hormone polyadenylation signal. Upon transfection into HEK 293T cells, Sec-gD was efficiently secreted into the culture supernatant **(Fig. 2C)**.

**Fig. 1.**
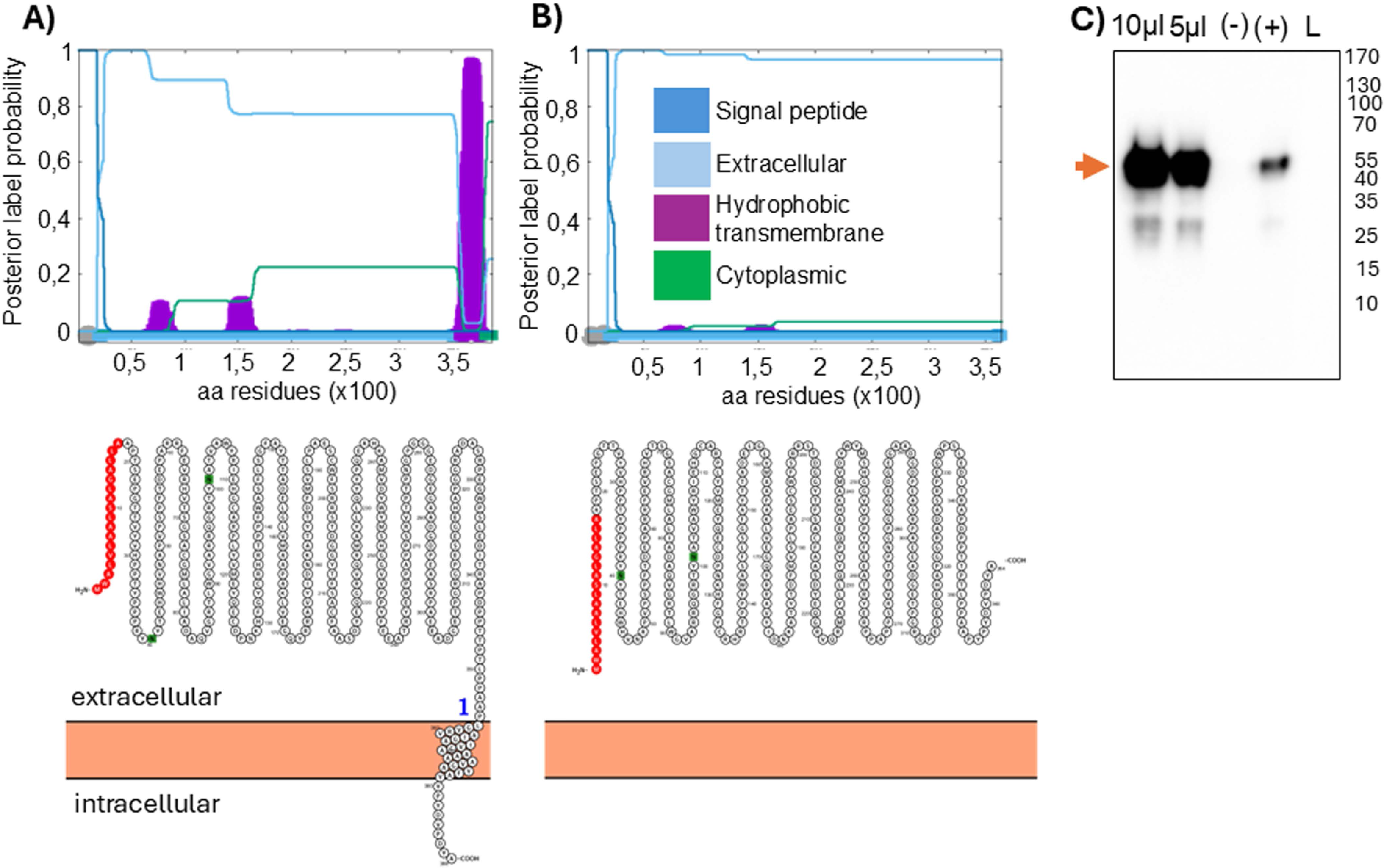
CpHV-1 sec-gD analysis and expression. Server output prediction of transmembrane topology and signal peptides from the amino acid sequence of gD (A) and sec-gD proteins (B) respectively. The plot shows the posterior probabilities of cytoplasmic (green)/extracellular(azure)/TM helix (purple)/signal peptide (blue) by calculating the total probability that a residue belongs to a helix, cytoplasmic, or non-cytoplasmic summed over all possible paths through the model. **C)** Western immunoblotting of sec-gD coming from transfected HEK 293T cells supernatant. Lanes were loaded with 10 and 5 ul of serum free medium transfected HEK 293T cells supernatant. Negative control was established pEGFPC-1 serum free medium transfected HEK 293T cells supernatant (*-*), whereas negative control was established pCMV-E2-HA serum free medium transfected HEK 293T cells supernatant (*-*).

**Fig. 2.**
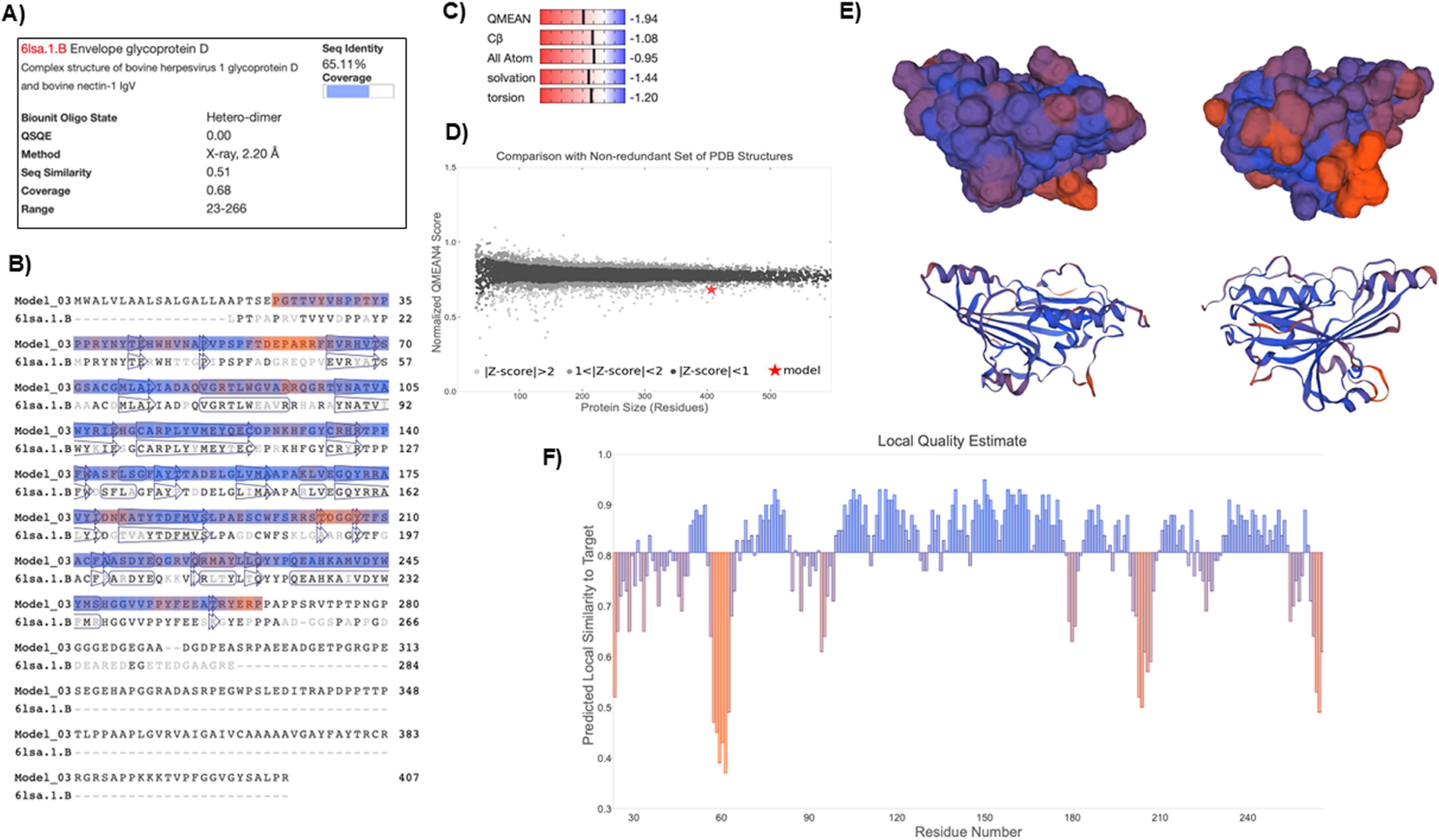
Template-Based Modeling of CpHV-1 gD Using SWISS-MODEL. The 3D structure of CpHV-1 glycoprotein D (gD) extracellular domain was predicted (residues 23–266) using the crystal structure of BoHV-1 gD (PDB ID: 6lsa.1.B) as a template, **(A)** Template details including sequence identity (65.11%) and coverage (68%) **(B)** Sequence alignment of the predicted CpHV-1 gD model with the BoHV-1 gD template, highlighting aligned residues and conserved secondary structures (Arrows - β-strands and cylinders - α-helices) **(C)** Global quality estimates using QMEAN scoring, indicating overall reliability of the predicted model. **(D)** Z-score plot comparing the model to a non-redundant set of PDB structure with similar size **(E)** 3D structure of the CpHV-1 gD computed model on two different orientations. **(F)** Local quality estimates per residue, with predicted local similarity to the template. Residues are colored based on predicted structural reliability (**blue** for well-conserved and confidently modeled regions, **orange** for areas of lower structural confidence).

### Sec-gD structure analysis

While structural data for gD proteins from other alpha herpesviruses are available, no such information currently exists for CpHV-1. To address this gap, the secondary structure of CpHV-1 gD was predicted using two computational tools. GOR IV (accessed on 28 May 2025) estimated the protein to consist of 61.43% random coil, 20.15% alpha-helix, and 18.43% extended strand. Similarly, PSIPRED 4.0 (accessed on 28 May 2025) predicted 71.79% random coil, 19.27% alpha- helix, and 8.94% extended strand **(Supplementary File 2)**. To model the 3D structure of CpHV-1 gD, template-based modeling was performed using the SWISS-MODEL web server (accessed on 28 May 2025). Among 172 matching templates, 10 were selected based on sequence coverage (0.53– 0.68), identity (43.32%–70.27%), and GMQE scores (0.39–0.51), all derived from previously resolved alpha herpesvirus gD structures **(Supplementary File 3)**. The final model was built using the bovine herpesvirus 1 gD structure (PDB ID: 6lsa.1.B), which shares 65.11% sequence identity and 68% coverage with CpHV-1 gD. The resulting model spans residues 23 to 266, encompassing the majority of the extracellular domain **(Fig. 2A)**. Model quality assessment yielded a GMQE of 0.51, a QMEANDisCo Global score of 0.81 ± 0.05 and a QMEAN Z-score of -1.94 **(Fig. 2C)**, indicating that the predicted structure is reasonably accurate and falls within the range of experimentally determined proteins of similar size **(Fig. 2D)**. Structural visualization and local quality estimation **(Figs. 2E and 2F)** showed high confidence in most surface-exposed regions, with lower reliability observed in loop regions around residues ∼55–65 and ∼200–210. To further refine the model, GalaxyRefine was employed. Both the initial and refined models were evaluated using the SWISS-MODEL structure assessment tool (accessed on 28 May 2025). Ramachandran plot analysis confirmed structural validity, with the refined model showing improved accuracy: 98.35% of residues were in favored regions with 0.00% outliers, compared to 95.04% favored and 0.83% outliers in the crude model **(Supplementary** Figure 4).

### Sec-gD exhibits conserved antigenic traits

The gD glycoprotein plays a crucial role in the attachment and entry of CpHV-1 into host cells (Keuser et al., 2006). Additionally, CpHV-1 gD is essential for eliciting neutralizing antibodies and conferring protection against experimental CpHV-1 infection in goats (Tempesta et al., 2000; Donofrio et al., 2013). Protein glycosylation can significantly influence antibody function, particularly affecting antigen recognition and binding affinity. This is especially relevant for conformational antibodies, which depend on the three-dimensional structure of the antigen and are therefore sensitive to glycosylation-induced changes (Tremain et al., 2023). The gD is a glycosylated protein (Keuser et al., 2006), including its secreted form (Sec-gD), which contains two predicted N-glycosylation sites at amino acid positions 40 and 101 **(Fig. 3A and B)**. These sites were experimentally confirmed through PNGase treatment and western blot analysis **(Fig. 3C**). Since structural prediction and analysis defined a certain degree of authenticity respect to other alpha herpesvirus glycoprotein D, it was of interest to ensure that Sec-gD retained its antigenic properties—despite modifications such as removal of the transmembrane domain and addition of a 9-amino-acid tag—a reverse serum neutralization assay was conducted **(Fig. 3D)**. As shown in **Fig. 3E**, preincubation of neutralizing sera with Sec-gD abolished their neutralizing activity, allowing CpHV-1 to infect and destroy the cell monolayer.

**Fig. 3.**
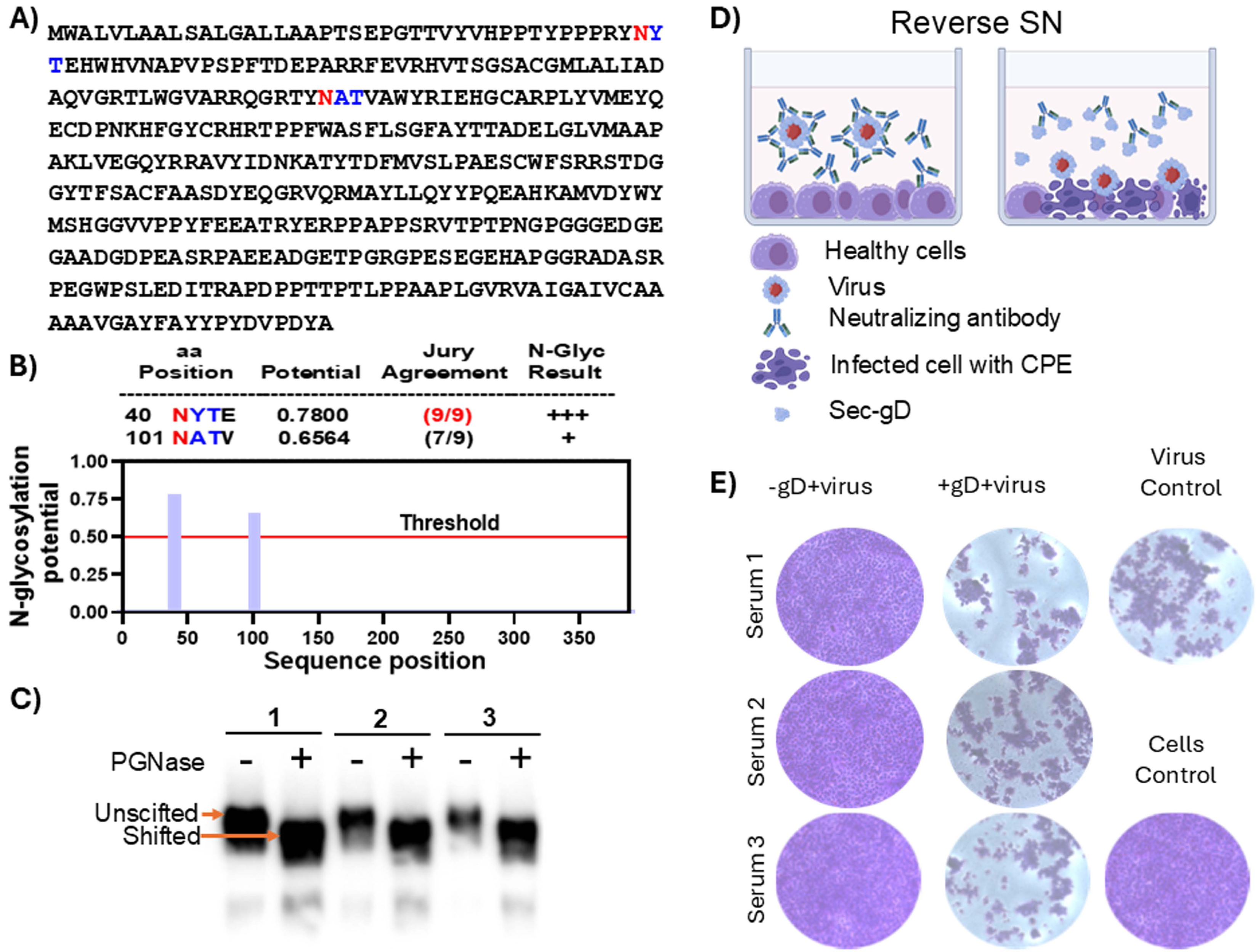
Sec-gD antigenic traits. **A)** sec-gD aminoacids sequence where N-linked glycosilation sites as predicted by NetGlyc 1.0. **B)** Potentially glycosylated Asparagines (N) are in red and flanked by consensus amino acids in blue. The value crossing the default threshold of 0.5, represents a predicted glycosylated site (as long as it occurs in the required sequence Asn-Xaa-Ser/Thr without Proline at Xaa). The ’potential’ score is the averaged output of nine neural networks. For further information, the jury agreement column indicates how many of the nine networks support the prediction. **C)** Western immuno blotting of sec-gD protein treated (+) or untreated (-) with PGNase F. The assay was repeated 3 times (1, 2, and 3) with different amount of protein to have a better resolution in terms of molecular size difference between glycosilated (Unshifted) and deglycosilated (Shifted) sec-gD. **D)** Reverse serum neutralization (SN) assay to test the authentic antigenic characteristics of sec-gD. Cartoon explaining the rational of the assay, where CapHV-1 neutralizing sera preincubated with medium containing sec-gD, are blocked by sec-gD interaction and allowing the virus to induce CPE. **E)** The quantitative assay performed in a 96-multiwell plate, 3 sera (1, 2, 3, and 4) containing neutralizing antibodies against CapHV-1 were tested at the dilutions of 1/10 in the presence of sec-gD (+sec-gD) and in the absence of sec-gD (−secgD). Virus Control was established in the absence of sera and sec-gD (Virus Control) and a Cells Control with cells without virus, sera and sec-gD (Cell Control). Crystal violet staining allows macroscopic (not shown) and microscopic evaluation of the integrity (violet wells) of the cell monolayer.

### Sec-gD allows the development of an indirect ELISA for detection of CpHV-1 infected goats

Sec-gD was used to coat 96-well ELISA plates, and ELISA assays were conducted using serial dilutions of individual serum samples. These included 10 sera from animals that tested negative by serum neutralization (SN) prior to experimental infection, and 10 sera collected from the same animals 42 days post-CpHV-1 infection, which tested SN-positive, clinically symptomatic and virologically shedding as monitored by clinical score from day 0 to day 11 post infection **(data not shown)**. The resulting dilution curves **(Fig. 4A)** and the corresponding average areas under the curves **(Fig. 4B)** allowed for clear discrimination between positive and negative sera. The serum dilution yielding the highest ratio between the average signal of positive and negative sera (P/N) was determined to be 1:40 **(Fig. 4C)**. ROC analysis **(Fig. 4D)** confirmed that 1:40 could represent the optimal serum dilution. Indeed, as shown by the area under the curve (AUC) values reported in Table 1, both the 1:40 and 1:80 dilutions gave an AUC value of 1. However, the maximum Youden index was obtained with different cut-off points: 0.199 for 1:40 and 0.127 for 1:80 **(Supplementary File 5)**. Therefore, the dilution with the higher cut-off was chosen, since for a diagnostic purpose a higher threshold is generally more robust against inaccurate readings at low absorbance level. This helps to prevent false positives resulting from background noise or potential weak cross-reactivity.

**Fig. 4.**
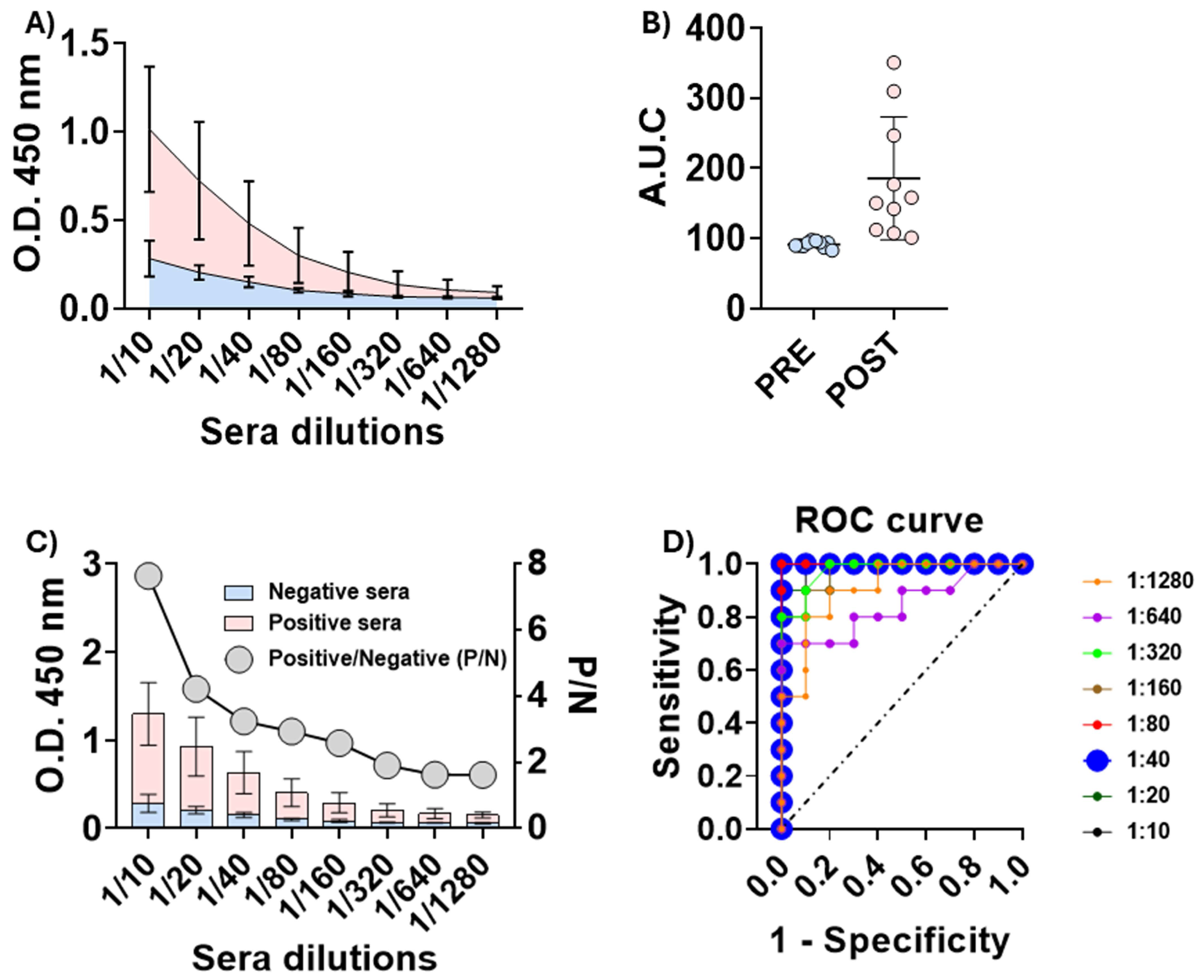
Reactivity of positive and negative sera toward sec-gD. **A)** Average reactivity of positive (pink) and negative (azure) sera at different dilutions. **B)** Data from the same experiment as in (A), but plotted as area under the curve (AUC) to obtain a better quantitative impression. Statistical analyses were performed using an unpaired two-tailed Student’s t-test in GraphPad Prism (P>0.0001). **C)** Optimization of sera dilution, defined as the dilution yielding the highest ratio between the average signal of positive and negative sera (P/N). D) ROC analysis to determine the cut-off value of the test.

**Table 1.** Area under the curve (AUC) values obtained at different dilutions of the tested sera. Sera dilutions with the best AUC are highlighted in bold.

| Serum dilution | AUC | Standard error | P value | 95% confidence interval |  |
| --- | --- | --- | --- | --- | --- |
|  |  |  |  | Lower limit | Upper limit |
| 1:10 | 0.990 | 0.016 | <0.001 | 0.958 | 1.022 |
| 1:20 | 0.980 | 0.026 | <0.001 | 0.929 | 1.031 |
| **1:40** | **1.000** | **0.000** | **<0.001** | **1.000** | **1.000** |
| **1:80** | **1.000** | **0.000** | **<0.001** | **1.000** | **1.000** |
| 1:160 | 0.970 | 0.032 | <0.001 | 0.907 | 1.033 |
| 1:320 | 0.975 | 0.029 | <0.001 | 0.919 | 1.031 |
| 1:640 | 0.845 | 0.092 | <0.001 | 0.665 | 1.025 |
| 1:1280 | 0.910 | 0.065 | <0.001 | 0.783 | 1.037 |

## Discussion

The glycoprotein D (gD) of several alpha herpesvirus plays a key role in the viral entry process, which typically occurs in two stages: initial attachment, where viral attachment proteins interact with receptors on the host cell surface, followed by membrane fusion that enables viral penetration (Zhong et al., 2023). Antibodies generated against gD can neutralize the virus, and the presence of these neutralizing antibodies in vaccinated animals is considered a more reliable indicator of vaccine efficacy compared to cell-mediated immunity (Georgopoulos and James, 2024). Despite the known molecular characteristics of CpHV-1 gD, no studies have yet explored the potential of using gD or its secreted form as a viable antigen for diagnostics and vaccine development. In this study, the secreted form of CpHV-1 glycoprotein D (Sec-gD) was successfully obtained and characterized.

The in silico-derived structure of Sec-gD displays features consistent with those of gD proteins from other alpha herpesviruses, including its glycosylation pattern. Biochemical analyses confirmed that Sec-gD is properly glycosylated. SDS-PAGE and Western blot analyses revealed that Sec-gD exhibits a slightly higher molecular weight than predicted based on its amino acid sequence, a discrepancy that was resolved upon treatment with PNGase F, indicating the presence of N-linked glycans. Glycans often dominate the surface of viral glycoproteins, making the viral glycome a key factor in shaping the antigenicity and immunogenicity. At one end of the spectrum, glycans can form essential components of epitopes recognized by neutralizing antibodies, thus playing a critical role in the design of effective immunogens. Conversely, the success of peptide-based and bacterially expressed protein vaccines demonstrates that viral glycosylation is not always essential. Nevertheless, native-like glycosylation can reflect proper protein folding and the presence of conformational epitopes. Moreover, strategic modifications beyond native glycan mimicry—such as altering glycosylation site occupancy or glycan processing—may enhance the immunogenicity and protective efficacy of vaccine antigens (Newby et al., 2024). Given that proper protein folding and glycosylation can indicate the presence of conformational epitopes involved in the induction and recognition of neutralizing antibodies, this was validated through a reverse neutralization assay (Newby et al., 2024). In this test, Sec-gD effectively inhibited the CpHV-1 neutralizing activity of sera from CpHV-1-infected animals, confirming its role in antibody recognition. For corroborating the antigenic properties of CpHV-1 Sec-gD expressed in mammalian cells, an indirect ELISA test was developed. The ELISA test demonstrated complete accuracy in distinguishing sera from infected and non-infected animals.

This study utilized a secreted form of the gD protein (Sec-gD), expressed in mammalian cells. The Sec-gD protein demonstrated excellent immunogenicity, highlighting its potential not only for diagnostic applications but also as a subunit vaccine capable of inducing strong neutralizing antibody responses. Its performance underscores the importance of developing effective immunogens for successful vaccine strategies. Previous research identified the CpHV-1 gD protein as a promising vaccine candidate due to its critical role in viral attachment and entry into host cells. Recent findings confirm that this glycoprotein elicits potent neutralizing antibody responses, reinforcing its value as a key target for vaccine development. Animal studies further support this, showing that gD-based vaccines can confer protective immunity. A recombinant BoHV-4-based vector vaccine expressing the full-length CpHV-1 gD has recently been developed and shown to provide robust protection against lethal CpHV-1 infection in goats. The Sec-gD platform represents an evolution of this approach, offering greater flexibility. It can be adapted into various formats, such as a ferritin-based self-assembling nanoparticle displaying the Sec-gD head domain (Sec-gD- ferritin), or as nucleic acid vaccines (DNA or mRNA). These formats offer several advantages, including rapid development, low-cost manufacturing, scalability, stability at lower temperatures, and suitability for deployment in outbreak-prone regions. Notably, DNA or mRNA-based Sec-gD vaccines are expected to elicit high levels of IFN-γ from T cells and induce strong gD-specific IgG antibody responses. Moreover, Sec-gD can be engineered into different structural configurations— monomeric, multimeric, or chimeric subunits—and stabilized in either fusion or post-fusion forms, enhancing its versatility as a vaccine component.

In conclusion, the promising results and proof-of-concept for using Sec-gD presented in this study lay the groundwork for developing and testing additional derivatives as prototype diagnostic tools and vaccines. These could target other significant animal pathogens—and potentially even human pathogens—given that CpHV-1 serves as a valuable large animal model for studying varicella-zoster virus in humans.

## Data Availability Statement

All data are available in this paper and Supplementary Materials, which can be retrieved by clicking the dedicated link.

## Acknowledgments

This study and the APC were funded by “Finanziamento dell’Unione Europea-Next generation EU- PRIN-2022/PNRR, grant number 2022M2HP84”. The funders had no role in study design, data collection and interpretation, or the decision to submit the work for publication.

The authors declare no conflicts of interest.

